# Protective effects of central leptin on whole-body energy homeostasis upon acute olanzapine exposure

**DOI:** 10.1101/2025.04.11.648424

**Authors:** Roshanak Asgariroozbehani, Raghunath Singh, Sally Wu, Ali Sajid Imami, Abdul-rizaq Hamoud, Sri Mahavir Agarwal, Bradley J. Baranowski, Stewart Jeromson, Ashley Bernardo, Thomas D. Prevot, David C. Wright, Adria Giacca, Robert E. Mccullumsmith, Sandra Pereira, Margaret K. Hahn

**Affiliations:** Centre for Addiction and Mental Health, Toronto, ON, Canada; Institute of Medical Sciences, University of Toronto, Toronto, ON, Canada; Department of Neurosciences and Psychiatry, University of Toledo, Toledo, OH, USA; Department of Psychiatry, University of Toronto, Toronto, ON, Canada; Banting & Best Diabetes Centre, Temerty Faculty of Medicine, University of Toronto, Toronto, ON, Canada; BC Children’s Hospital Research Institute, Vancouver, BC, Canada; School of Kinesiology, Faculty of Education, University of British Columbia, Vancouver, Canada; Faculty of Land and Food Systems, University of British Columbia, Vancouver, BC, Canada; Department of Pharmacology & Toxicology, University of Toronto, Toronto, ON, Canada; Department of Physiology, University of Toronto, Toronto, ON, Canada; ProMedica, Neuroscience Institute, Toledo, OH, USA

**Keywords:** antipsychotics, olanzapine, energy homeostasis, metabolism, hypothalamus, kinome arrays

## Abstract

Second-generation antipsychotic use is associated with severe metabolic side effects such as obesity and type 2 diabetes. Leptin is a hormone that is secreted by adipose tissue, and it acts on the brain to decrease body weight by reducing food intake and stimulating energy expenditure. Leptin also improves glucose and lipid metabolism. We examined the short-term impact of olanzapine, a commonly used second-generation antipsychotic, on the central leptin-mediated regulation of energy balance, lipid metabolism, and hypothalamic kinase activity. Male Sprague Dawley rats were given an acute intracerebroventricular (ICV, 3^rd^ ventricle) injection of either leptin or vehicle, combined with subcutaneous olanzapine or vehicle. As expected, ICV leptin decreased food intake and importantly, olanzapine did not block this effect. Administration of leptin, olanzapine, or their combination reduced the average respiratory exchange ratio (RER) during the light cycle, which indicates that fat oxidation was increased. In the dark cycle, leptin decreased the average RER regardless of olanzapine administration, and in the presence of leptin, olanzapine did not affect the average RER. Leptin did not alter the olanzapine-induced increase in serum triglyceride concentrations. Olanzapine and central leptin treatment differentially activated hypothalamic kinases. In conclusion, regulation of food intake and fuel preference by central leptin is intact following acute olanzapine administration.

## INTRODUCTION

Antipsychotics (APs) are the mainstay treatment for schizophrenia spectrum disorders and are also prescribed off-label in the management of other mental health conditions such as bipolar disorder, major depression disorder, autism spectrum disorder, Tourette’s Syndrome, and insomnia (1–4). Despite efficacy in alleviating psychiatric symptoms and safety over first-generation APs (e.g., chlorpromazine, and haloperidol) in inducing extrapyramidal side effects, use of second-generation APs (e.g., clozapine, olanzapine (OLA), risperidone, etc.) is associated with a high incidence of metabolic side effects including weight gain, hyperglycemia, type 2 diabetes, and dyslipidemia (3, 5, 6). These metabolic disturbances contribute to increased rates of cardiovascular disease, which is a significant cause of premature death in schizophrenia patients (7, 8). AP-induced metabolic disturbances can occur independently of changes in body weight (7, 8). Several mechanisms have been explored to delineate AP-associated metabolic perturbations, including disrupted hypothalamic appetite control (9, 10), and impaired nutrient sensing (11, 12), insulin and leptin resistance (13–15), adiposity and inflammation (16, 17), and changes in gut microbiota (18). Prolonged AP exposure in rodents leads to weight gain and adiposity changes, along with increased food intake (19, 20). OLA exposure increases free fatty acid uptake and promotes lipogenesis in white adipose tissues (21, 22). The effect of OLA on lipolysis appears to be time-dependent, with stimulation in the short-term and inhibition in the long-term (5, 23). Accordingly, acute OLA treatment induces a shift in fuel preference towards fat oxidation rather than carbohydrate oxidation, as indicated by a reduction in respiratory exchange ratio (RER) (22).

Leptin is a hormone that is secreted mainly from white adipose tissue (24). It decreases food intake and body weight mainly by acting on the brain, especially the hypothalamus (25–27). Leptin improves glucose metabolism and increases energy expenditure and fat oxidation; it also inhibits lipogenesis and stimulates lipolysis and fat oxidation in white adipose tissue (24, 28–30). These effects can be induced by intracerebroventricular (ICV) or intrahypothalamic administration of leptin, indicating the importance of central leptin in the regulation of whole-body metabolism (30–33).

Several studies have reported significantly higher leptin concentrations in multiple episode (chronic) schizophrenia patients compared to healthy controls (34–36). This has been largely attributed to chronic AP treatment and subsequent weight gain, particularly in those treated with second-generation APs; notably, OLA, clozapine, and quetiapine were associated with the most substantial increases in leptin concentrations, emphasizing their higher metabolic risk compared to other APs (34–36). Leptin is secreted in proportion to white adipose tissue mass (24), but Zhao et al. (15) found that OLA and risperidone cause a rapid increase (within 3 days) in circulating leptin levels even before inducing weight gain in female mice fed high-fat diet. Furthermore, leptin neutralization (i.e., treatment with leptin antibody) prevented hyperleptinemia, weight gain, increased food intake, glucose intolerance, hyperinsulinemia, and inflammation (systemic, adipose tissue, and hypothalamic) in mice chronically treated with risperidone and OLA (15). We previously reported that OLA injected subcutaneously upregulates inflammatory pathways in the hypothalamus within two hours of administration (12, 37). In the current study, we used an acute model to determine if OLA disrupts the effects of central leptin on energy balance, including food intake and energy expenditure, and glucose as well as lipid metabolism, in the absence of AP-induced body weight gain. Furthermore, we used a protein kinase activity assay and associated pathway analysis to determine the effects of leptin and OLA on the kinome in the hypothalamus.

## MATERIALS AND METHODS

### Animals

Animal use protocol was approved by the Animal Care Committee (ACC) at the Centre for Addiction and Mental Health (CAMH) and followed the Canadian Council on Animal Care (CCAC) guidelines. Adult male Sprague Dawley rats (300–400 g, Charles River, Saint-Constant, QC, Canada) were maintained on a 12 h light/dark with ad libitum access to food and water. We have previously used the same animal model, namely male Sprague Dawley rats, to demonstrate that olanzapine impairs the ability of central insulin to reduce food intake (38). Animals were pair-housed and acclimatized to the facility.

### Surgical Procedures

After approximately 1 week of acclimatization in the animal facility, rats underwent intracerebroventricular (ICV) cannulation surgeries according to the previously described method (12, 39). Briefly, a cannula (HRS Scientific Guide 38172, 22 gauge, cut 9 mm below pedestal) was implanted into the third ventricle to target the hypothalamus, following stereotaxic coordinates: anterior–posterior −2.5 mm, and dorsal–ventral −8.0 mm. Two adjutant stainless steel screws (McCray Optical Supply Inc., Scarborough, ON, Canada) were fixed, and dental cement used to hold the cannula. Rats were allowed to recover for 1 week after the ICV surgery, including 3 days of postoperative care, and singly housed.

### Experimental design

Rats were randomly assigned to experimental groups based on ICV - Leptin (Lep) or vehicle (Veh); and subcutaneous (SC) - Olanzapine (OLA) or Veh treatments. Lyophilized leptin (Recombinant rat leptin, Sigma-Aldrich) was dissolved in sterile saline (NaCl 0.9%), while OLA (Toronto Research Chemicals, Toronto, ON, Canada) was dissolved in sterile saline using 10% glacial acetic acid (10µl/ml) for a final concentration of 0.1%. Across all three experiments, 3µl of Lep (1µg/µl) was administered ICV (40, 41) using Hamilton syringe, while OLA was injected subcutaneously at the dose of 2 mg/kg (42). Experimental groups were as follows: (i) Veh-Veh; (ii) Veh-OLA; (iii) Lep-Veh; (iv) Lep-OLA.

#### Experiment 1. Indirect calorimetry in metabolic cages

After 1 week of recovery from ICV surgery, rats (n=6-7 per group) were acclimated to powdered chow (LabDiet 5001) in the Comprehensive Laboratory Animal Monitoring System (CLAMS) indirect calorimetry cages, also known as metabolic cages (CLAMS; Oxymax, Columbus Instruments), for 3 days. The data collected during the first 24h is not included here because it was used to ensure baseline parameters were similar between rats. Rats were then randomly assigned to the experimental groups. On Day 2, at the start of the light cycle (8 AM) and dark cycle (8 PM), animals received two treatments: SC (Veh/Ola) followed by ICV (Veh/Lep) and were placed in the metabolic cages. Metabolic parameters including oxygen consumption (VO_2_), carbon dioxide production (VCO_2_), respiratory exchange rate (RER), food intake, heat expenditure, and locomotive activity were recorded every 10 minutes by CLAMS for 24 hours (38). Afterwards, the animals were placed back in their home cages. A schematic representation is given in Fig.1A.

**Figure 1.**
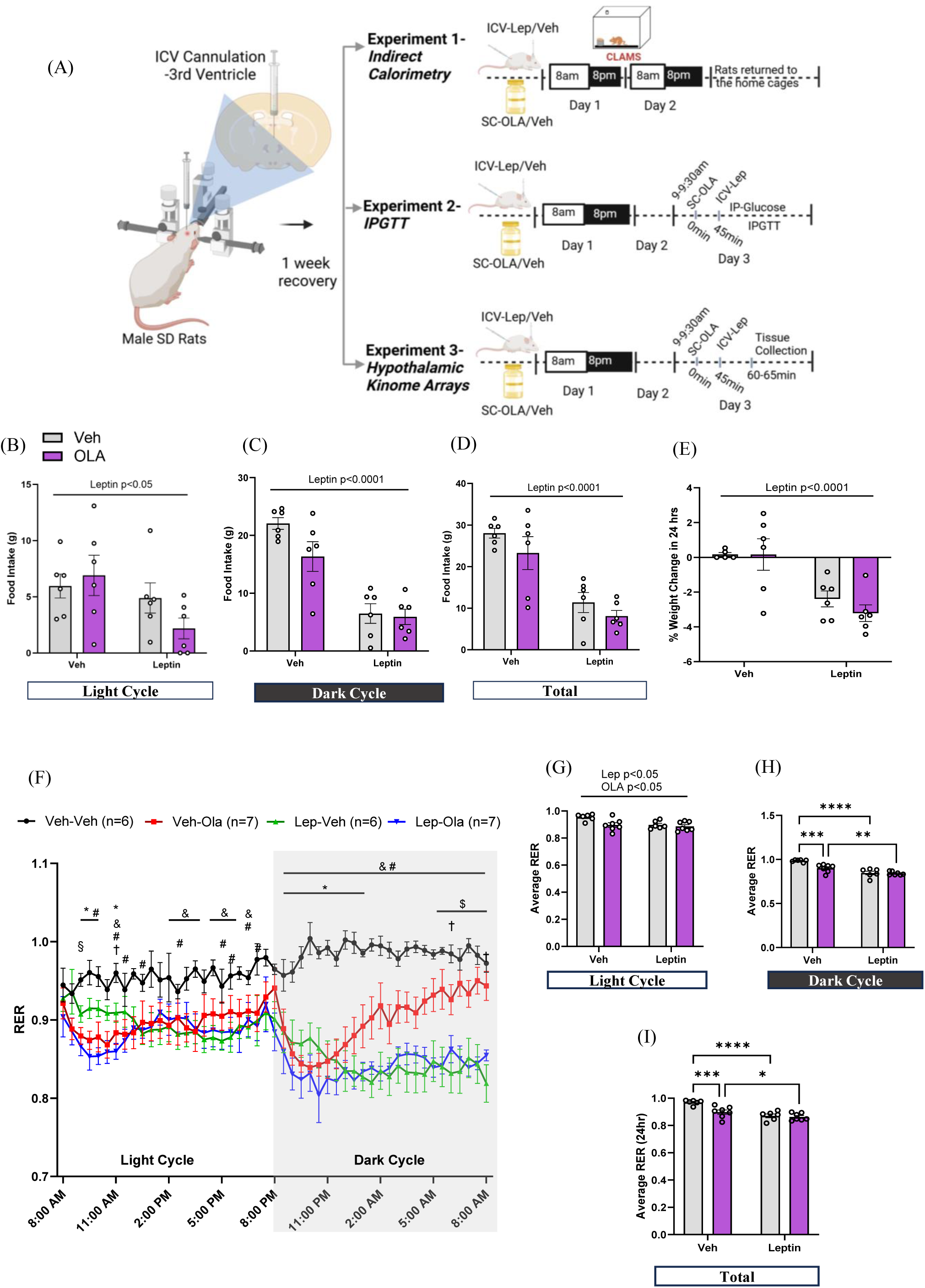
Effect of acute olanzapine and ICV-leptin treatment on food intake, fuel preference, and body weight. Rats (*N*= 6-7) were treated with ICV-leptin (Lep) or vehicle (Veh) along with SC-olanzapine (OLA) or vehicle (Veh) at the start of light and dark cycles; changes in metabolic parameters were recorded by indirect calorimetry cages in 10-min intervals. **(A)** Schematic representation of experimental design (figure made with BioRender). **(B-E)** Cumulative food intake in light and dark cycles, and percentage change in body weight in 24-hour duration of treatment. Percentage change in body weight was obtained from a second cohort of rats. **(F)** Line graph showing 24-hour RER starting from 08:00AM-08:00PM (light cycle), 08:00PM-08:00AM (dark cycle); timepoints shown are at every 30 min. * p<0.05, Veh-OLA vs. Veh-Veh; & p<0.05, Lep-Veh vs. Veh-Veh, # p<0.05, Lep-OLA vs. Veh-Veh; $ p<0.05, Veh-OLA vs. Lep-OLA; † p<0.05, Veh-OLA vs. Lep-Veh; § p<0.05, Lep-Veh vs. Lep-OLA. (**G-I)** Respiratory exchange ratio (RER), including average RER in light cycle, dark cycle and over 24h. ** p<0.01, *** p<0.001, and **** p<0.0001. Comprehensive Laboratory Animal Monitoring System-CLAMS, Leptin-Lep, Olanzapine-OLA, Vehicle-Veh.

#### Experiment 2. Intraperitoneal glucose tolerance test (IPGTT)

A separate cohort of rats (n=6-8 per group) was used for assessing effect of acute OLA and ICV Lep on glucose metabolism during the IPGTT. Like in the previous experiment, rats received two treatments: ICV (Veh/Lep) and SC (Veh/OLA) at the start of the light cycle (8-9 AM) and dark cycle (8-9 PM) on the day 1. On Day 2, rats were food restricted overnight. On Day 3, animals received respective treatments (same as on day 1) and then subjected to the IPGTT. A baseline glucose reading was acquired from the rats (t=0) before administering the SC (Veh/OLA) treatment. After 45 minutes of SC treatment, ICV (Veh/Lep) treatment was administered and this was immediately followed by an intraperitoneal injection of 2 g/kg dextrose (50%; Pfizer, Canada). Blood glucose was measured at 15, 30, 60, 90, and 120 minutes, from the tail tip using a glucometer (Accu-Chek). A schematic representation of the methodology is illustrated in Fig. 1A.

#### Experiment 3. Acute treatment for ex vivo molecular analyses

To investigate the molecular mechanism underlying the acute effects of OLA and ICV leptin on energy homeostasis, and glucose and lipid metabolism, another cohort of rats (n=6-8 per group) was used. Similar to the previous experiments, rats received treatments twice: ICV (Veh/Lep) and SC (Veh/OLA) at the start of the light cycle (8-9 AM) and dark cycle (8-9 PM) on the day 1. On the second day, animals were food restricted overnight. On Day 3 (at 9 AM), rats received SC injection of Veh/OLA and after 45 minutes, ICV (Veh/Lep) treatment was administered. Approximately 20 min post ICV treatments, rats were euthanized under anesthesia (schematic representation provided in Fig.1A). Blood following decapitation was collected and serum was collected. Epididymal white adipose tissue (eWAT) and hypothalamus were harvested, snap frozen, and kept frozen for molecular assays.

### Biochemical and Molecular experiments

#### Serum analysis

Serum concentration of free fatty acids (FFAs) (Fujifilm Wako Pure Chemical Corporation,), triglycerides (Triglyceride Colorimetric Assay Kit, Cayman Chemical), and glycerol (Glycerol Colorimetric Assay Kit, Cayman Chemical) were measured following the manufacturer instructions. Serum insulin was determined using an ultrasensitive rat insulin ELISA kit (Mercodia), and glucose was measured using Analox GM9 Glucose analyzer. Fasting serum glucose and insulin values were used to calculate Homeostatic Model Assessment for Insulin Resistance (HOMA-IR) for estimation of insulin resistance.

#### Immunoblotting

Protein expression of enzymes involved in lipolysis, namely hormone-sensitive lipase (HSL) phosphorylated (pHSL (s563), Cell Signaling 4139S) and total HSL (tHSL, Invitrogen PA1-16966) and total adipose triglyceride lipase (ATGL, Cell Signaling 2138S), was assessed in eWAT using immunoblotting as described previously (43). Phosphorylation of HSL is a marker of its activity. Briefly, eWAT was homogenized, protein extracted and quantified, and membranes were incubated overnight at 4°C with gentle rocking in primary antibodies diluted (1:1000) with TBST and 5% BSA. The following morning, membranes were briefly washed in TBST and incubated for 1 h at room temperature with horseradish peroxidase-conjugated secondary antibodies (1:2000). Proteins of interest were expressed relative to an internal ponceau loading control from the same respective membrane as the protein of interest. Immunoblots were quantified using ImageJ 1.53e software, National Institute of Health, USA.

#### Hypothalamic Kinome Array

Hypothalamic samples were prepared and analyzed by kinome array as previously described (12, 37), except that in the current study chips reporting tyrosine kinase activity were also used, and samples for each treatment group were pooled and run as technical replicates (n=3). Only kinases with a Z score ≥ ǀ1.5ǀ are included in this manuscript. Kinome array involves adding hypothalamic samples to chips with immobilized non-phosphorylated peptides: either serine/threonine kinase (STK) chips or phospho-tyrosine kinase (PTK) chips. These peptides can be phosphorylated by known serine/threonine or tyrosine kinases. Antibodies specific for phosphorylated peptides are then added to the chips and the binding of these phosphoantibodies to their target sequences on the chips generates a signal. A greater signal implies increased activity of the kinases that phosphorylate that peptide sequence and vice versa. Kinase families were identified using KRSA (kinome random sampling analyzer) (44–46) (Supplementary Excel file 1). To identify specific kinases in families with Z score ≥ ǀ1.5ǀ based on KRSA, PTM-SEA (posttranslational modifications-signature enrichment analysis) (47, 48) was used. In the current study, we focused on the following 3 group comparisons: Lep-Veh vs. Veh-Veh, Veh-OLA vs. Veh-Veh, and Lep-OLA vs. Lep-Veh. Pathway analysis used WikiPathway 2023 (Human) (Supplementary Excel files 2 and 3).

### Statistical analysis

All data are presented as means ± SEM and also individual data points. Significance was defined as p<0.05; n=6-8 rats were used in each group. Statistical analyses were performed using GraphPad PRISM 10. Repeated measure ANOVAs were used for analysis of RER and IPGTT results (main effects were group and time); if there was a main effect of group or a significant statistical interaction, Tukey’s post-hoc test was performed at each timepoint (i.e., for RER, 6 group comparisons were done at each of the 49 timepoints; for IPGTT, 6 group comparisons were done at each of the 6 timepoints). Food intake (light cycle, dark cycle, or total), VO2 (light or dark cycle), VCO2 (light or dark cycle), heat production (light or dark cycle), locomotion (light or dark cycle), and area under the curve for IPGTT were analyzed using a two-way ANOVA (2 main effects were ICV treatment and SC treatment). If the statistical interaction was significant for each of these parameters, then Fisher’s LSD test was used for the following 4 group comparisons: Lep-Veh vs. Veh-Veh, Veh-OLA vs. Veh-Veh, Lep-OLA vs. Lep-Veh, Lep-OLA vs. Veh-OLA.

## RESULTS

### ICV-leptin suppresses food intake and body weight even in the presence of OLA

Leptin treatment alone or in combination with OLA (i.e., Lep-Veh and Lep-OLA) significantly reduced food intake during the light cycle, dark cycle, and both (p<0.05, p<0.0001, and p<0.0001, respectively, Fig. 1B-D). Reduced food intake was associated with body weight reduction in leptin-treated groups (p<0.0001; Fig. 1E).

### RER decreases following individual and combined treatment with OLA and ICV-leptin

A reduction in RER indicates increased fat oxidation relative to oxidation of the other macronutrients (carbohydrates and proteins). Acute treatment with OLA (Veh-OLA) resulted in rapid decrease in RER which was most prominent 2-3 hours post-treatment compared to the vehicle (Veh-Veh) group (Fig. 1F). A rapid reduction in RER was observed in the dark cycle following SC-OLA treatment, but RER gradually increased towards the end of the dark cycle in the Veh-OLA group. Treatment with ICV-leptin alone (Lep-Veh) resulted in a gradual reduction in RER in the light and dark cycles. Co-treatment with ICV-leptin and SC-OLA (Lep-OLA) decreased RER in the light cycle, which became even more prominent in the dark cycle following re-administration of treatments. In Fig. 1 G, H, and I, the average RER during the light cycle, dark cycle, or over 24 h are shown and the results recapitulate the individual data points shown in Fig. 1F. In the light cycle, both leptin and OLA decreased average RER (p<0.05; Fig. 1G). In the dark cycle, leptin decreased average RER, regardless of SC treatment (p<0.0001, and p<0.001), OLA decreased RER only in the absence of leptin (p<0.01), and among groups treated with leptin, OLA did not significantly alter RER. Similar results were obtained for the average 24h RER (Fig. 1I).

OLA increased oxygen consumption (VO_2_) during the light cycle (p<0.01). During the dark cycle, VO_2_ was decreased by leptin (p<0.01, Fig. 2A and 2B). Carbon dioxide production (VCO_2_) tended to be increased by OLA in the light cycle (p=0.0588, Fig. 2C). In the dark cycle, VCO_2_ was decreased by leptin, regardless of treatment (p<0.0001 and p<0.01, Fig. 2D), and OLA diminished VCO_2_ only in the absence of leptin (p<0.05, Fig. 2D). Although physical activity (ambulance and rearing counts) was similar between groups during the light cycle, OLA alone (Veh-OLA vs. Veh-Veh) and leptin alone (Lep-Veh vs. Veh-Veh) decreased (p<0.001 and p<0.0001) physical activity in the dark cycle (Fig. 2E-H). Heat production, which is indicative of energy expenditure, was increased by OLA in the light cycle (p<0.001; Fig. 2I), and decreased by leptin in the dark cycle (p<0.05; Fig. 2J).

**Figure 2.**
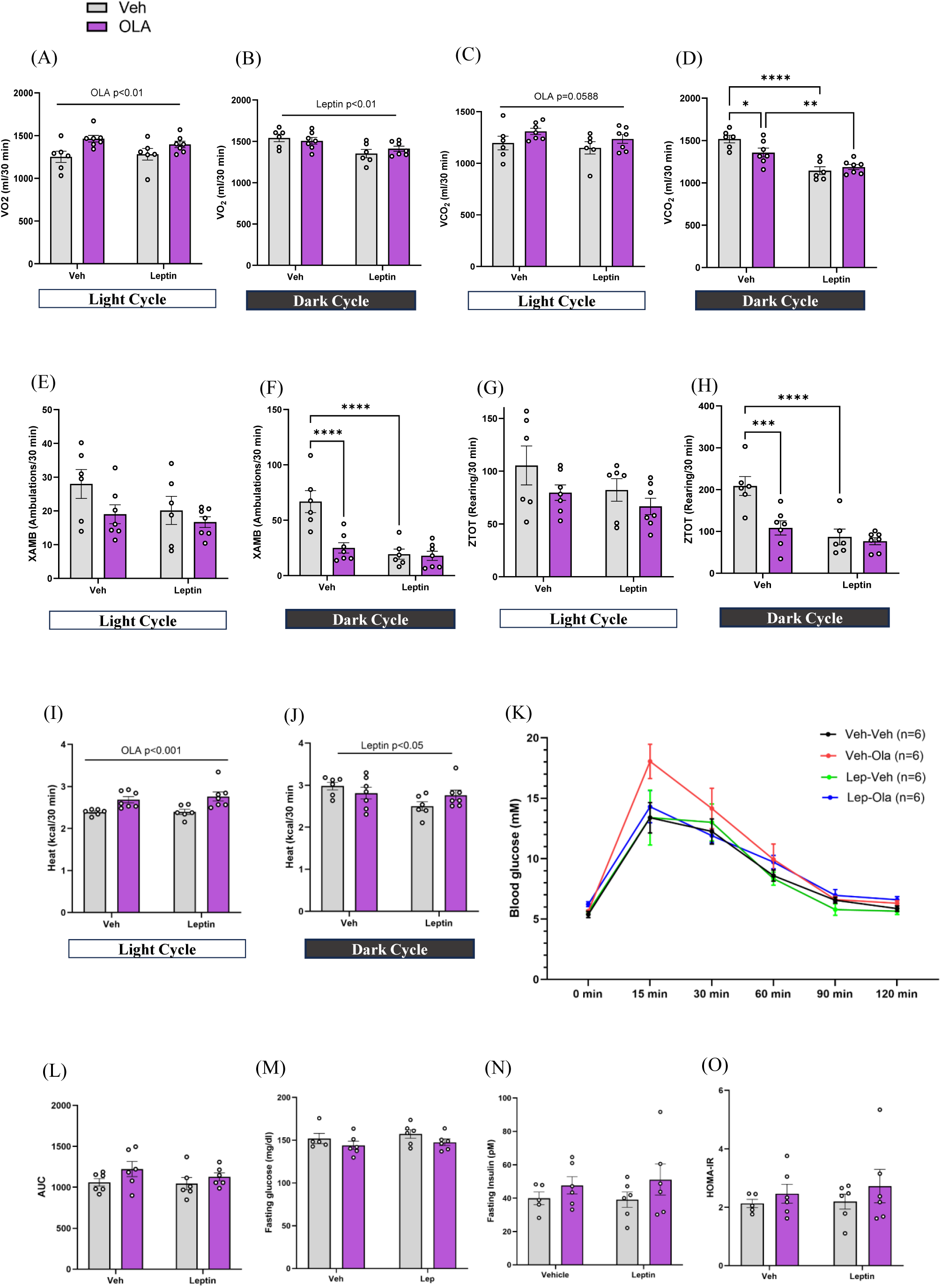
Effect of acute olanzapine and ICV-leptin treatment on VO_2_, VCO_2_, locomotion and heat production in metabolic cages, and on glucose metabolism. Rats (*N*= 6-7) were treated with ICV-leptin (Lep) or vehicle (Veh) along with SC-olanzapine (OLA) or vehicle (Veh) at the start of light and dark cycles; changes in metabolic parameters were recorded by indirect calorimetry cages in 10-min intervals. **(A-D)** Volume of oxygen (VO_2_), and carbon dioxide (VCO_2_) in light and dark cycle. (E-H) Locomotor activity in light and dark cycle measured as total ambulation and rearing counts. **(I-J)** Heat production. Overnight food restricted rats (*N*=6 per group) were subjected to an intraperitoneal glucose tolerance test (IPGTT) and another cohort of rats (*N*=5-7 per group) was used for serum collection. **(K-L)** Effect of acute SC-olanzapine and ICV-Leptin exposure on glucose concentrations during IPGTT and respective area under the curve (AUC). There is a time effect (p<0.0001) in **K**. **(M)** Fasting serum glucose, (**B)** Fasting serum insulin, **(C)** Homeostatic Model Assessment for Insulin Resistance (HOMA-IR). * p<0.05, ** p<0.01, *** p<0.001, and **** p<0.0001. Leptin-Lep, Olanzapine-OLA, Vehicle-Veh.

### Neither OLA nor leptin affect glucose tolerance

There were no differences in glucose tolerance between the 4 groups following an IPGTT (Fig. 2K and 2L). There was, however, a trend for the IPGTT AUC to be increased by OLA (main effect, p=0.085) (Fig. 2L). Furthermore, leptin and OLA did not alter fasting serum glucose and insulin concentrations (Fig. 2M-N). Accordingly, HOMA-IR, an index of insulin sensitivity that depends on fasting glucose and insulin concentrations, was similar between groups (Fig. 2O).

### ICV-leptin and OLA affect certain aspects of lipid metabolism

Treatment with OLA increased serum triglyceride concentrations (p<0.05, Fig. 3A). OLA itself (Veh-OLA vs. Veh-Veh) and leptin itself (Lep-Veh vs. Veh-Veh) increase serum FFA concentrations (p<0.01 and p<0.05, respectively, Fig. 3B). Interestingly, among groups treated with OLA, leptin decreased serum FFA concentrations (Veh-OLA vs. Lep-OLA, p<0.05). Serum glycerol concentrations were similar across groups (Fig. 3C). Since increased serum FFA concentrations may indicate increased adipose tissue lipolysis, we performed immunoblotting to assess lipolytic markers (ATGL, pHSL, tHSL and pHSL/tHSL) in eWAT. Acute leptin and OLA treatment did not change the protein expression of ATGL, pHSL, and tHSL (Fig. 3D-G). However, regardless of ICV treatment, OLA decreased the ratio of pHSL to tHSL, which suggests decreased lipolysis (p<0.05, Fig. 3H).

**Figure 3.**
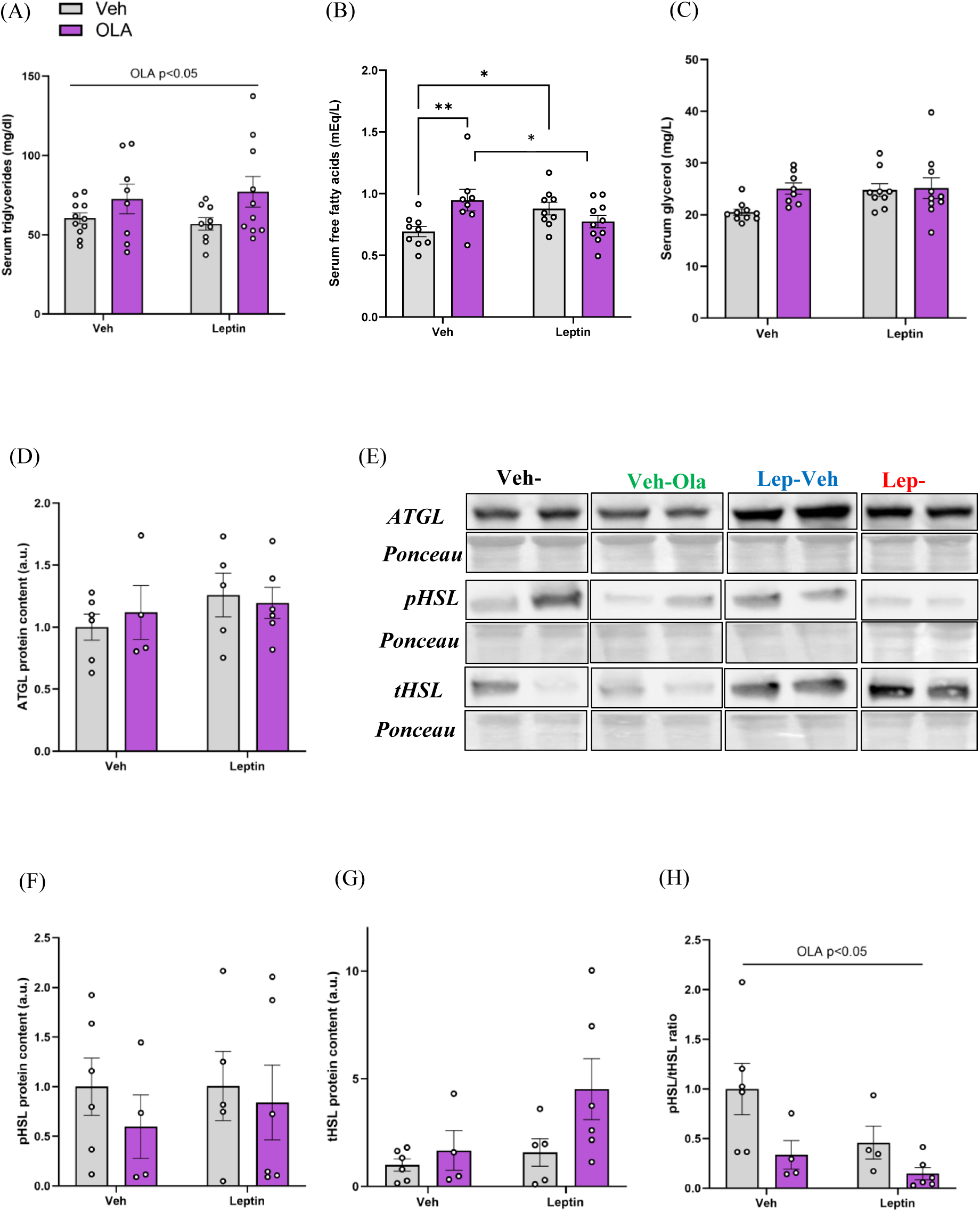
Effect of olanzapine and ICV-leptin on lipid metabolism. Overnight food restricted rats were used for serum and tissue (eWAT and hypothalamus) collection. (**A-C)** Serum concentrations of triglycerides, free fatty acids, and glycerol (N= 9-11 in each group). * p<0.05 and ** p<0.01. **(D-H)** Protein expression of ATGL, pHSL, tHSL, and ratio of pHSL/tHSL in eWAT and representative images of the gels from immunoblotting (N= 4-6 in each group). Leptin-Lep, Olanzapine-OLA, Vehicle-Veh, ATGL-adipose triglyceride lipase, p/tHSL-phosphorylated/total hormone-sensitive lipase.

### Acute ICV-leptin and OLA treatments differentially alter activities of hypothalamic protein kinase activities

The overlap in kinase activities between the 3 comparisons is shown in Fig. 4A. In Figures 4 (B, C, D) and 5 (B, C, D), the dots represent peptides targeted by the kinases stated on the x-axis. The red dots are relative signal intensity of peptides that are below a 0.85 fold change (i.e., log2 fold change ≤ −0.2) or above a 1.15 fold change (i.e., log2 fold change ≥ 0.2). The location of the majority of red dots for a given kinase determines how its activity was affected; therefore, if a given kinase has the majority of the red dots above a 1.15 fold change, then the activity of that kinase was increased. Since in our study ICV-leptin has a beneficial effect on key parameters independent of OLA treatment or prevented the untoward effects of OLA, our interpretation of kinase activity results focused on comparisons that could explain this effect. Specifically, we focused on: 1) kinases whose activity was altered by leptin (i.e., in the Lep-Veh vs. Veh-Veh comparison), or by OLA itself (Veh-OLA vs. Veh-Veh), but was unaffected by OLA in the presence of leptin (i.e., in the Lep-OLA vs. Lep-Veh comparison), and 2) kinases whose activity was altered specifically in the Lep-OLA vs. Lep-Veh comparison, but was not affected by leptin itself (i.e., in the Lep-Veh vs. Veh-Veh comparison) or by OLA itself (i.e., Veh-OLA group vs. Veh-Veh group).

**Figure 4.**
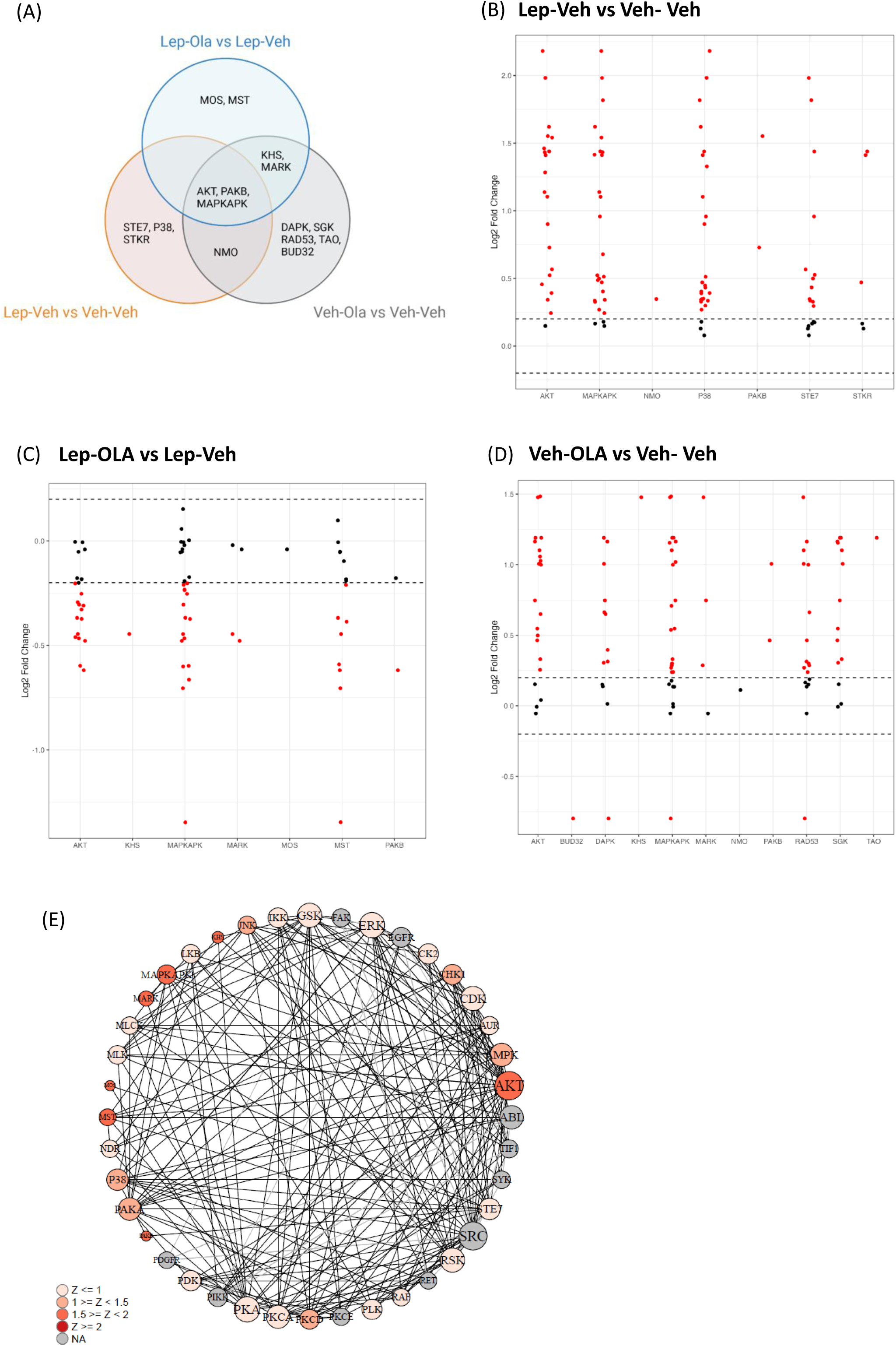
Effect of acute ICV leptin and olanzapine treatment on activity of hypothalamic serine/threonine kinases. **(A)** Venn diagram showing overlap in the kinase activity among all the group comparisons (Veh-Ola vs Veh-Veh, Lep-Veh vs Veh-Veh, and Lep-Veh vs Lep-Ola). (**B-D)** Log2 fold change in relative signal intensity of reporter peptides for serine/threonine kinase activity; kinases on the x-axis refer to kinase families for the group comparisons (n=3 in each group). In A-D, only kinases with a Z score ≥ ǀ1.5ǀ are included. (**E)** Kinase network model for comparison of Lep-Ola vs Lep-Veh, where the circle diameter increases with interaction number. Leptin-Lep, Olanzapine-OLA, Vehicle-Veh.

Results of serine/threonine kinase activity for each of the 3 comparisons are found in Fig. 4A-E. Fig. 4A shows overlap and differential kinase activities among all 3 comparisons; alterations in AKT, MAPKAPK, and PAKB activity are common in all 3 comparisons. Leptin increases the activity of various serine/threonine kinases (Fig. 4B), while the activity of serine/threonine kinases is decreased by OLA in the presence of leptin (Fig. 4C). OLA itself mostly increases the activity of serine/threonine kinases (Fig. 4D). Notably, p38 MAPK, STE7, and STKR activity is increased by leptin, but OLA does not alter this effect of leptin. The corresponding kinase network shows the prominent kinases that can be differentiated by the bigger circle, which represents higher interaction with other proteins (Fig. 4E). Analysis using PTM-SEA indicates that within the p38 MAPK family, it is the p38α isoform (MAPK14) that is altered (Supplementary Fig. 1). The overlap in tyrosine kinase activity between the 3 comparisons is in Fig 5A. Tyrosine kinase activity for each of the 3 comparisons is found in Fig. 5 B-D. Overall, leptin itself increases tyrosine kinase activity that largely remains when OLA is present.

**Figure 5.**
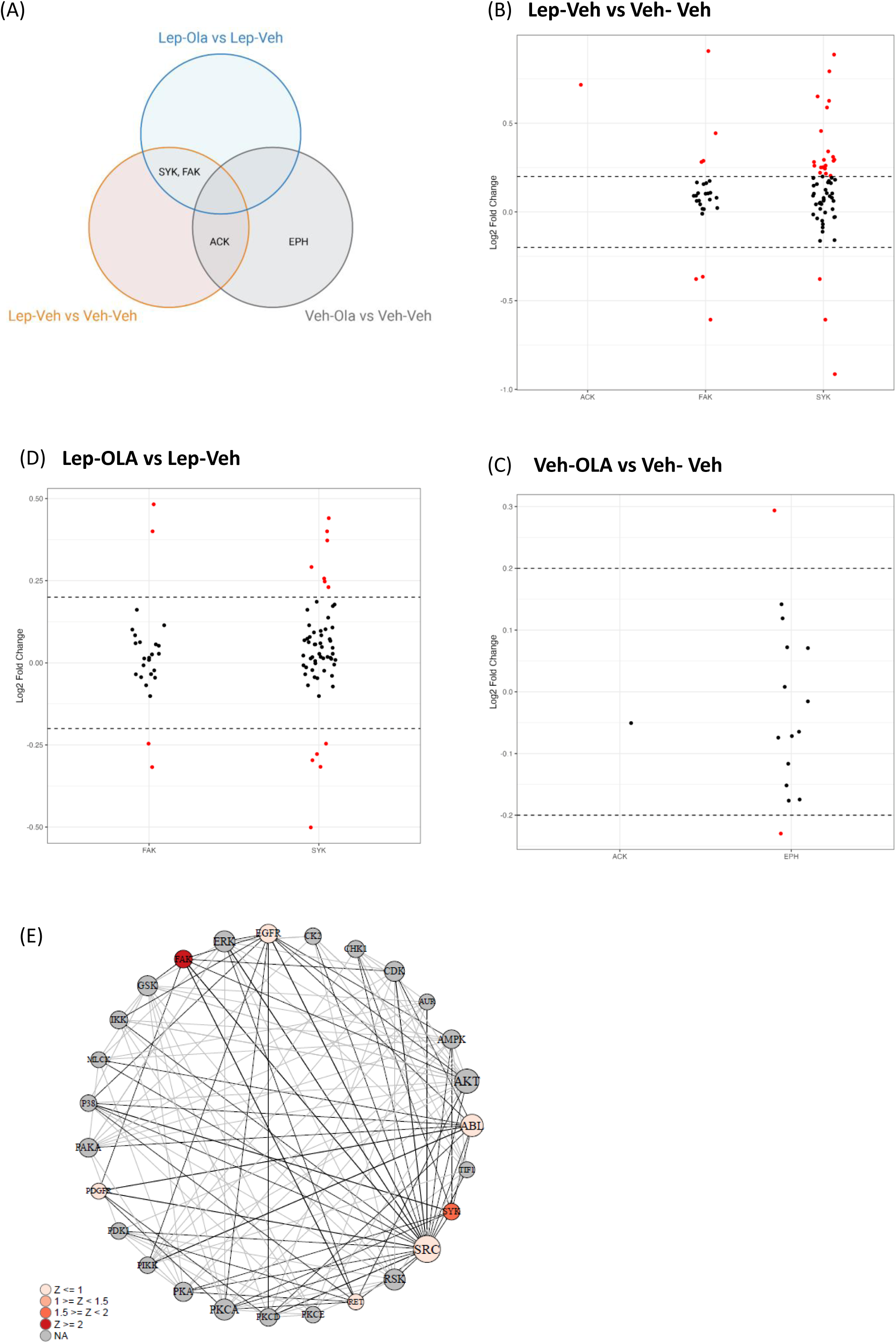
Effect of acute ICV leptin and olanzapine treatment on activity of hypothalamic tyrosine kinase activity. **(A)** Venn diagram showing overlap in the kinase activity among all the group comparisons (Lep-Veh vs Veh-Veh, and Lep-OLA vs Lep-Veh, Veh-OLA vs Veh-Veh). (**B-D)** Log2 fold change in relative signal intensity of peptides for tyrosine kinase activity; kinases on the x-axis and refer to kinase families for the group comparisons (n=3 in each group). In A-D, only kinases with a Z score ≥ ǀ1.5ǀ are included. (**E)** Kinase network model for comparison of Lep-OLA vs Lep-Veh, where the circle diameter increases with interaction number. Leptin-Lep, Olanzapine-OLA, Vehicle-Veh.

Pathway analysis based on kinome array results (STK and PTK chips) was also performed. Overall direction of change of pathways that were statistically significant (based on adjusted p value) is based on whether there was a net increase or decrease in the phosphorylation of peptides for each comparison. For the Lep-Veh vs. Veh-Veh comparison, there was a net increase in serine/threonine phosphorylation of peptides and in tyrosine phosphorylation of peptides (Supplementary Fig. 2 and 3). For the Lep-OLA vs. Lep-Veh comparison, there is a net decrease in serine/threonine phosphorylation of peptides, while the net change in tyrosine phosphorylation of peptides is close to zero, thereby making interpretation of this pathway analysis difficult (Supplementary Fig. 4 and 5). For the Veh-OLA vs. Veh-Veh comparison, there is a net increase in serine/threonine phosphorylation of peptides, while the net change in tyrosine phosphorylation of peptides is close to zero, thereby making interpretation of this pathway analysis difficult (Supplementary Fig. 6 and 7).

Keeping these phosphorylation results in mind, as expected, ICV leptin stimulated the leptin signaling pathway based on both STK and PTK chip results (Supplementary Excel files 2 and 3). Moreover, ICV leptin stimulated insulin signaling, brain-derived neurotrophic factor (BDNF) signaling pathway, p38 MAPK signaling pathway, and pro-inflammatory pathways. Somewhat surprisingly given our *in vivo* results, in the context of leptin (Lep-OLA vs. Lep-Veh), OLA impairs leptin, insulin, and BDNF signaling pathways. However, there is an attenuation of inflammatory pathways in the presence of OLA. Interestingly, like leptin by itself, OLA alone stimulates leptin, insulin, and BDNF signaling pathways. It is unclear if this is due to OLA-induced alterations in circulating hormones, for example increased plasma insulin. OLA alone also increases inflammatory pathways in the hypothalamus.

## DISCUSSION

The key finding of our study is that acute subcutaneous OLA treatment did not impair the inhibition of food intake and decrease in body weight caused by central leptin. Certain aspects of lipid metabolism, such as OLA-induced increase in serum triglyceride concentrations, were not improved by central leptin administration.

We infused leptin into the 3^rd^ ventricle to target the hypothalamus. Leptin action in the arcuate nucleus of the hypothalamus is well-established to reduce food intake (49, 50). Accordingly, we found that ICV leptin reduced food intake and body weight. Surprisingly, hypophagia and decreased body weight caused by ICV leptin was observed despite OLA administration. In contrast, we have previously found that acute OLA administration overrides central insulin’s ability to reduce food intake in the dark cycle (when rodents are most active and eat the most) (38). Therefore, while acute OLA administration at a clinically relevant dose impairs central insulin’s hypophagic effects, it leaves central leptin-induced hypophagia intact. Acute OLA administration in the absence of ICV leptin did not increase food intake, in agreement with our previous study (38).

Chronic AP treatment causes hyperleptinemia, which in turn can cause a proinflammatory state and leptin resistance (15, 51). Zhao et al. (15) reported that a leptin-neutralizing antibody, which diminishes AP-induced hyperleptinemia, prevents weight gain caused by prolonged AP treatment. The authors also noted that AP-induced hyperleptinemia occurs within 3 days of AP administration and precedes weight gain. Our results support these findings because acute central leptin administration, likely before hyperleptinemia occurs, maintains energy balance in the context of OLA administration. It is important to note that central leptin signalling inhibits leptin production and secretion by white adipose tissue (52) and therefore, our results suggest that maintaining central leptin signalling intact may delay the onset of hyperleptinemia and the ensuing weight gain caused by APs.

Although OLA increased energy expenditure (heat) during the light cycle, regardless of ICV treatment, leptin decreased energy expenditure in the dark cycle, regardless of OLA treatment. The decrease in energy expenditure caused by leptin in the dark cycle may be secondary to the decrease in body weight over 24h resulting from leptin administration (Fig. 1E). The alterations in energy expenditure caused by leptin in the dark cycle may also be due to changes in physical activity, since reduced physical activity can decrease energy expenditure. In our study, leptin or OLA alone reduced physical activity during the dark cycle. A mechanism through which leptin can diminish locomotor activity is the inhibition of ventral tegmental area (VTA) dopaminergic neurons (53). In our study, this would likely result from leptin activating its receptors in the lateral hypothalamus (LH) and activating a LH-VTA circuit (53) because it is unclear if leptin injected in the 3^rd^ ventricle can travel to the VTA to directly engage its receptors there. OLA reduces locomotor activity because of its well-documented sedative effects (42, 54). A higher dose of OLA was not used in this study to prevent possible disruptions in normal feeding behavior.

RER values are closer to 1.0 in the dark cycle, indicating predominant use of carbohydrates. A reduction in RER suggests an increase in the use (oxidation) of fat and it is typically associated with the fasting state (light cycle in rodents and dark cycle in humans). Metabolic flexibility refers to the body’s capacity to adjust fuel oxidation depending on the metabolic state (55, 56). In our study, acute OLA treatment induced a rapid reduction in RER in both light and dark cycles, suggesting increased fat oxidation. Previous studies have demonstrated similar findings suggesting that OLA impairs metabolic flexibility by preventing the body from switching to carbohydrate oxidation (22, 38, 57). However, OLA-induced reductions in RER may be more of marker of metabolic disturbance, rather than a cause of metabolic impairment because ICV-leptin also reduced RER in both light and dark cycles, consistent with previous studies (36, 55). This result is reflective of leptin’s physiological effects including stimulation of fat oxidation, along with increase in lipolysis and inhibition of lipogenesis (24). It may also be secondary to the leptin-induced reductions in food intake (58). In groups that received ICV leptin, OLA did not further decrease average RER in the dark cycle. While the increase in fat oxidation caused by central leptin does not appear to impair carbohydrate metabolism, OLA may increase fat oxidation because it impairs carbohydrate metabolism (22). Moreover, OLA-induced impairments in carbohydrate metabolism are associated with depletion of Krebs cycle intermediates, which in turn are generated from amino acids (22). Indeed, OLA affects tissue levels of amino acids (59). It is also important to note that while a decrease in RER from 1.0 may be indicative of increased fat oxidation, it may also indicate an elevation in protein oxidation since the RER for fat oxidation is ~0.7 and for protein oxidation it is ~0.8. More research needs to be conducted to determine the mechanisms through which OLA affects RER.

Leptin has a catabolic effect on lipids, but OLA promotes lipid accretion (5, 29). While central leptin inhibits adipose tissue lipogenesis and simulates lipolysis, OLA stimulates adipogenesis; OLA’s effects on lipolysis appear to be time-dependent, stimulating it acutely and inhibiting it in the long-term (5, 21, 31, 33). In the current study, we cannot conclude convincingly whether leptin and olanzapine affect lipolysis because circulating markers of lipid metabolism, which depend on various tissues, do not agree with markers of lipolysis in white adipose tissue. We did not find group differences in the protein expression of ATGL and total HSL in white adipose tissue (eWAT). Nevertheless, we found that OLA inhibits a marker of HSL activity (ratio of phosphorylated to total HSL) in white adipose tissue, consistent with OLA’s anti-lipolytic effects reported in the literature. Yet, OLA increased circulating free fatty acid concentrations, but only in the absence of leptin. This acute effect of OLA has been reported by others (23). Leptin increased circulating free fatty acid concentrations (Lep-Veh vs. Veh-Veh), which may suggest increased lipolysis and/or lipoprotein lipase activity (29). However, in the presence of OLA, leptin decreased serum free fatty acid concentrations (Veh-OLA vs. Lep-OLA). OLA increased serum triglyceride concentrations, which have been reported in the literature and may be the result of OLA’s effects on lipid metabolism in the liver (5). Central leptin did not block the increase in serum triglycerides or the decrease in the marker of white adipose tissue lipolysis (pHSL/tHSL) resulting from OLA administration, which suggests that leptin’s protective effects may not extend to all aspects of lipid metabolism.

Acute and chronic treatment with OLA is associated with hyperglycemia, insulin resistance and impaired glucose homeostasis. These effects may occur through both central and peripheral mechanisms (11). Leptin improves glucose metabolism and insulin sensitivity (60, 61). In our study, neither OLA nor leptin were found to alter glucose tolerance or an index of insulin sensitivity (HOMA-IR). An oral glucose tolerance test (OGTT) stimulates the secretion of incretin hormones like glucagon-like peptide-1 (GLP-1) (62) and since GLP-1 receptor agonists improve glucose metabolism in rodents treated with olanzapine (63), it is possible that an OGTT could have uncovered differences in glucose tolerance. Furthermore, a more sensitive and complex technique for assessment of insulin sensitivity *in vivo*, such as the hyperinsulinemic euglycemic clamp, may have detected differences in insulin sensitivity.

The kinome array results suggest two mechanisms through which leptin action on food intake and body weight regulation remains intact in the presence of OLA *in vivo*. First, central inflammation causes hyperphagia (64) and we found that leptin has anti-inflammatory effects in the presence of OLA. However, OLA itself triggers hypothalamic inflammation, as found in the current study and in our previous study using RNA-seq results (37). OLA itself also causes ER stress in the hypothalamus (37) and ER stress can be upstream of inflammation. Therefore, in the current study, we propose that OLA, which is administered ~45 minutes before ICV leptin, causes hypothalamic ER stress and inflammation that is diminished upon administration of ICV leptin. Similar findings have been found by Gan et al. (65), where administration of leptin alleviated the ER stress and inflammation caused by a compound (tunicamycin). Our findings also reinforce the complex relationship between leptin and inflammation that has been documented in the literature because leptin itself stimulated inflammation. Hence, the context of leptin treatment is important. Despite OLA causing inflammation in our model, this may not translate to hyperphagia due to OLA’s sedative effects and the acuteness of the experiment. Second, leptin increases hypothalamic p38 MAPK activity and this is unaltered by OLA. It has been reported that leptin activates p38 MAPK, thereby promoting expression of neurotensin, which is a neurotransmitter that decreases food intake (66). Activation of p38 MAPK by leptin does not appear to be mediated by the tyrosine kinase JAK2.

The current study has some limitations. We chose a frequently used dosage of bolus ICV-leptin injection (67–69), but our study was acute. Similarly, acute dosing of OLA is not representative of its chronic, long-term, clinical use in patients. Future studies should also include female rats as second-generation APs can have different metabolic disturbances in male and female rodents (70–72). Although we found that neither OLA nor leptin affected glucose tolerance during an IPGTT, we cannot exclude the possibility that differences in glucose metabolism may be detected with more sensitive techniques.

In conclusion, we found that, acutely, central leptin reduced food intake even in the presence of OLA. Based on our study results and those of others (15), we propose that if hypothalamic leptin signalling is preserved within hours of initiating OLA treatment and before a rise in plasma leptin concentrations, which appears to take days, it may be possible to prevent or diminish OLA-induced weight gain. However, leptin may not be able to overcome certain aspects of lipid metabolism that are disturbed by OLA. More research is needed on the therapeutic potential of leptin in AP-induced metabolic dysfunction.

## AUTHOR CONTRIBUTIONS

**Asgariroozbehani Roshanak:** Conceptualization, Data curation, Writing original draft.

**Singh Raghunath:** Conceptualization, Data curation, Writing original draft, review & editing.

**Wu Sally:** Data curation.

**Hamel Laurie:** Conceptualization

**Prevot Thomas D**: Data curation, Formal analysis

**Bernardo Ashley:** Data curation, Formal analysis

**Agarwal Sri Mahavir:** Writing – review & editing.

**Baranowski Bradly:** Data curation.

**Jeromson Stewart:** Data curation.

**Wright David C.:** Funding acquisition, Writing – review & editing.

**Giacca Adria:** Writing – review & editing.

**Imami Ali Sajid**: Data curation, Formal analysis, Software.

**Hamoud Abdul-rizaq:** Data curation, Formal analysis, Software.

**Mccullumsmith Robert E.:** Funding acquisition, Supervision, Writing – review & editing.

**Pereira Sandra:** Data analysis, Writing original draft, Writing – review & editing.

**Hahn Margaret K:** Conceptualization, Funding acquisition, Project administration, Supervision, Writing – review & editing.

## ACKNOWLEDGEMENTS

MKH is supported by the CAMH and University of Toronto Meighen Family Research Chair. She is also supported by a research grant from Investigator-Initiated Studies Program of Merck Canada Inc. (MISP). D. H. Gales family charitable foundation, Banting and Best Diabetes Centre (BBDC), University of Toronto; Canadian Institutes of Health Research (CIHR); and CAMH discovery funds are gratefully acknowledged for providing postdoctoral fellowships to RS. REM is supported by funding from the National Institute of Mental Health (MH107487, MH121102) and National Institute of Aging (AG057598).

---

**Supplementary Figure 1.**
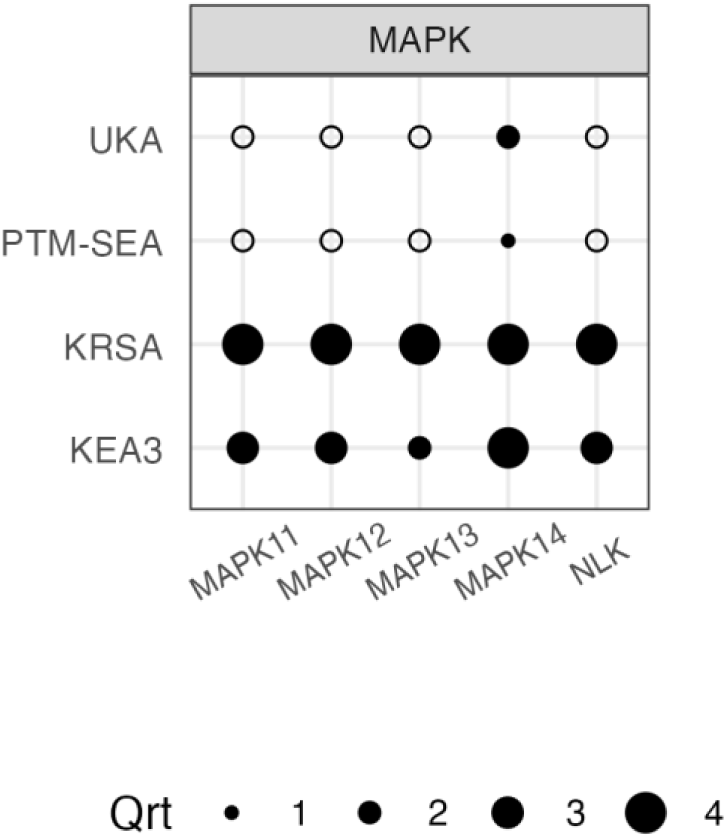
Different deconvolution methods to determine serine/threonine kinase activity in Lep-Yeh vs. Yeh-Yeh comparison. Diameter of black dots represent quartiles.

**Supplementary Figure 2.**
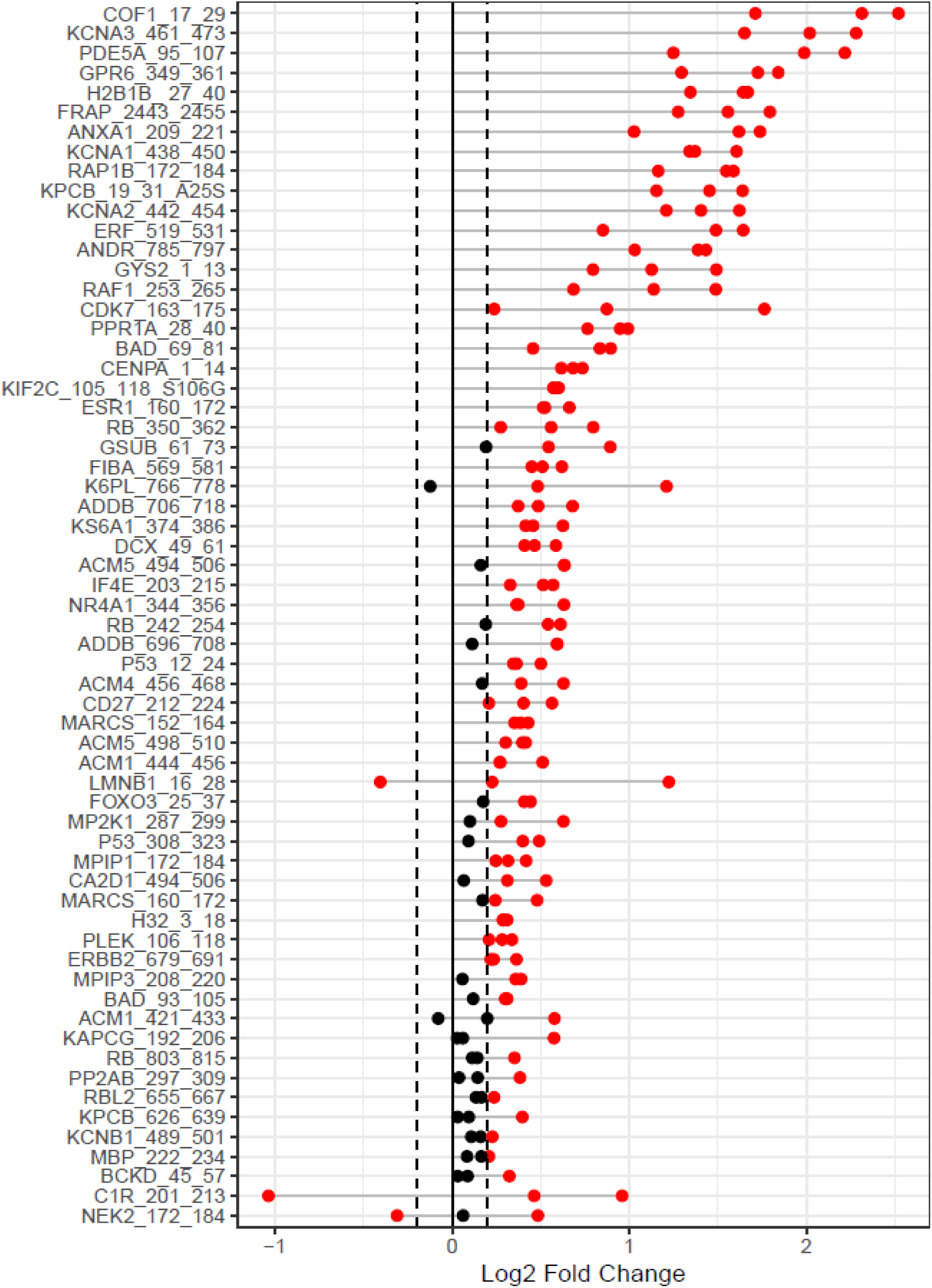
Waterfall Plot to show the distribution of change in peptide phosphorylation for STK, Lep-Yeh vs Yeh-Yeh.

**Supplementary Figure 3.**
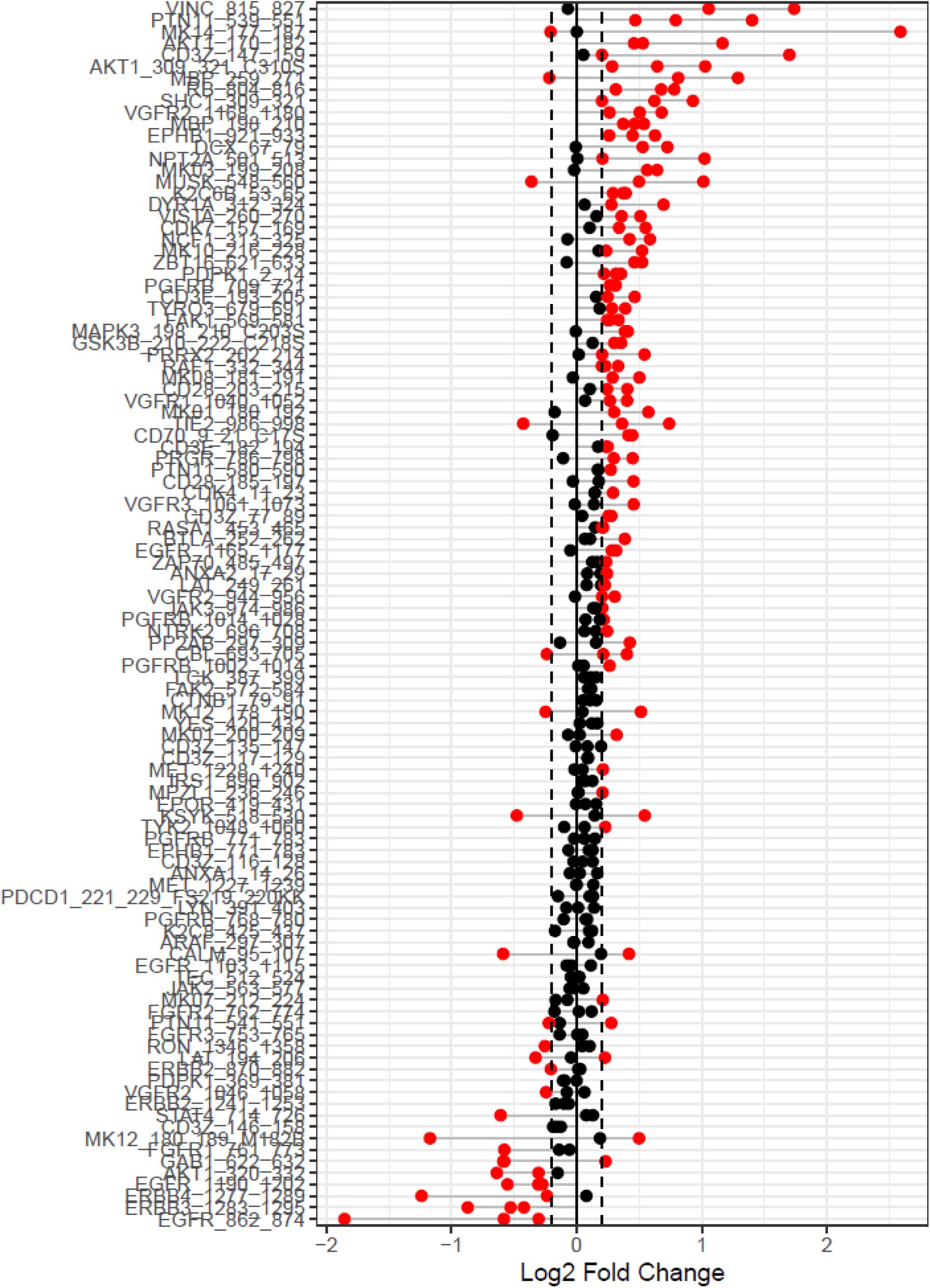
Waterfall Plot to show the distribution of change in peptide phosphorylation for PTK, Lep-Yeh vs Yeh-Yeh.

**Supplementary Figure 4.**
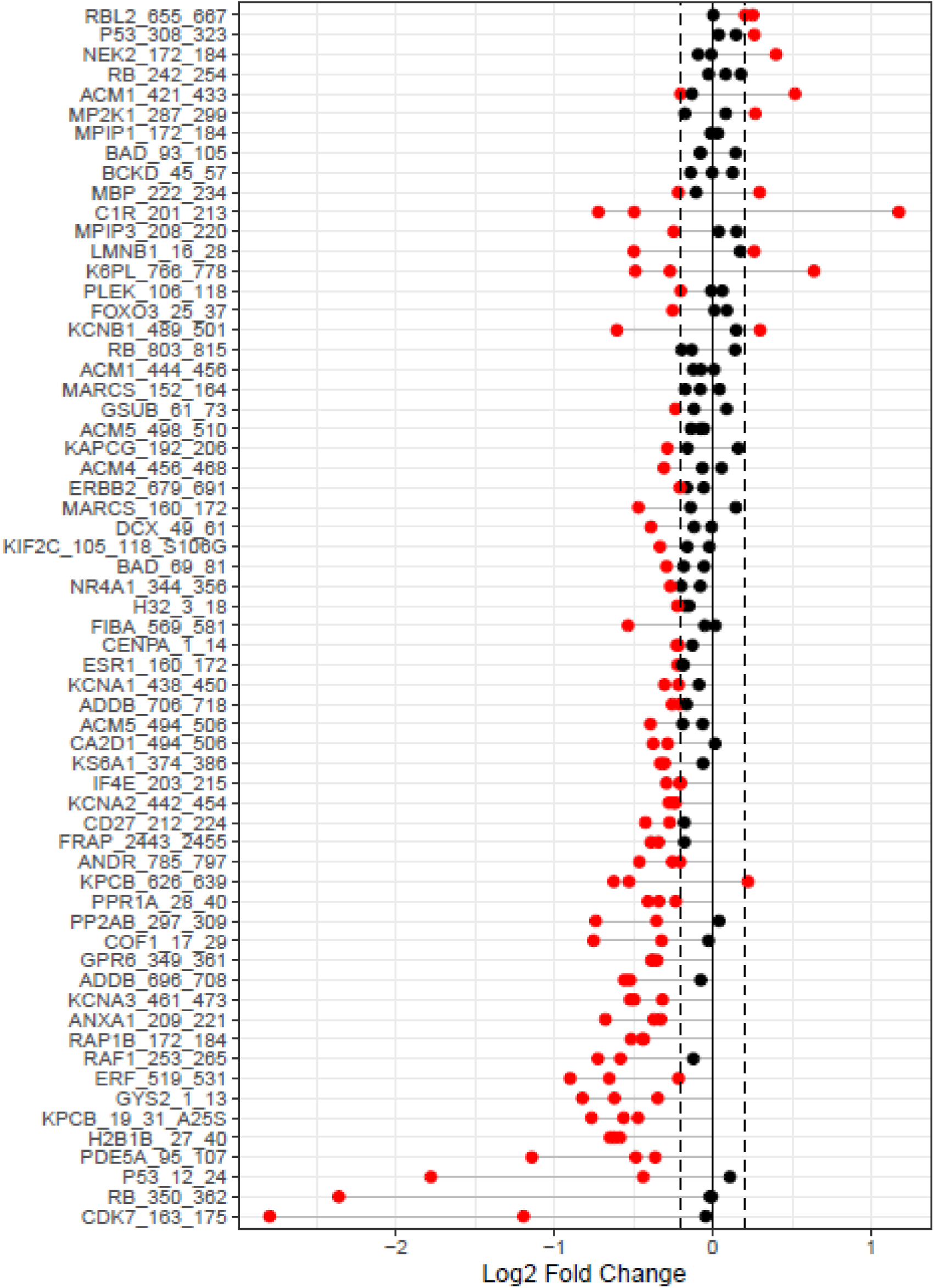
Waterfall Plot to show the distribution of change in peptide phosphorylation for STK, Lep-OLA vs Lep-Yeh.

**Supplementary Figure 5.**
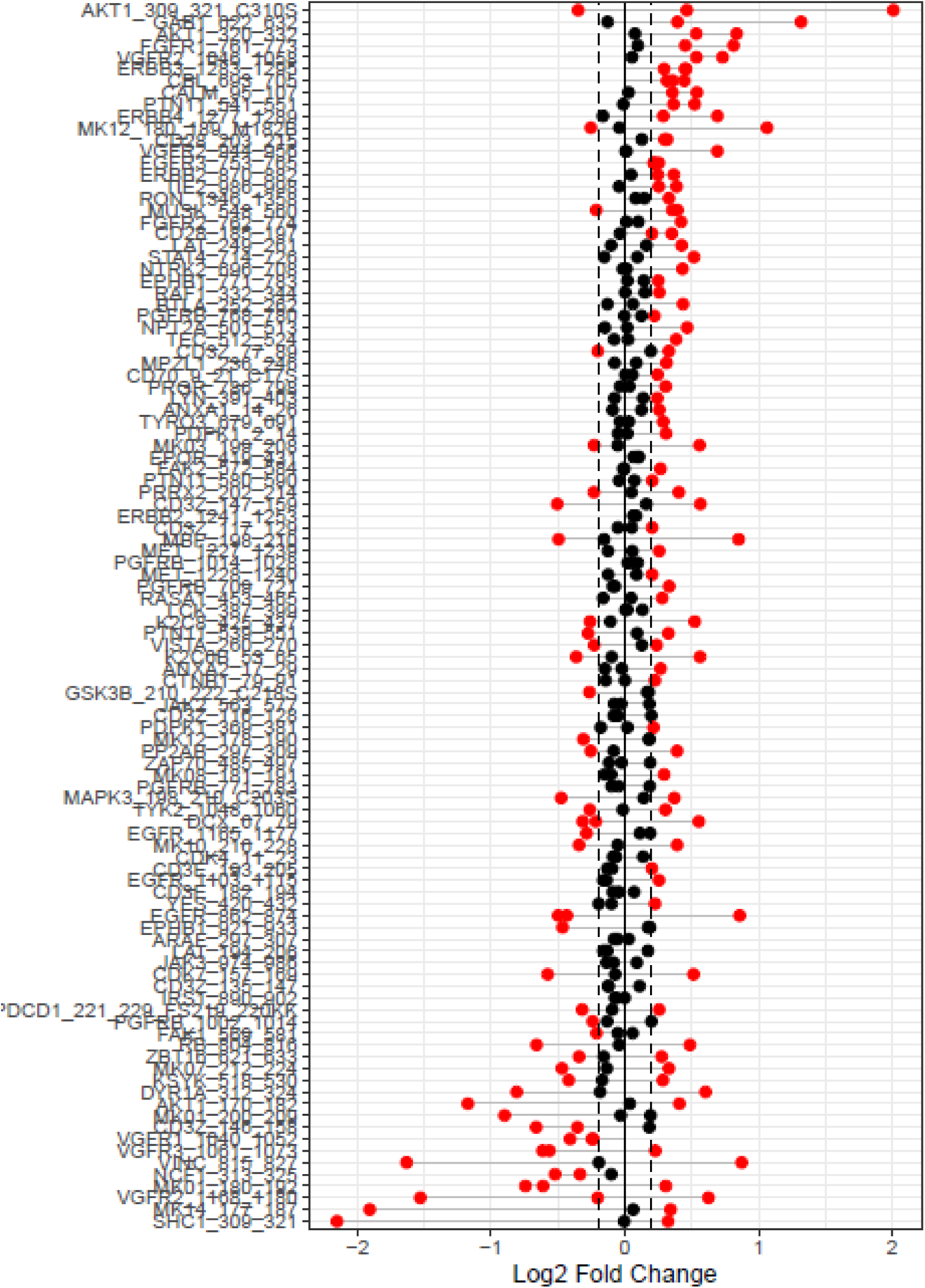
Waterfall Plot to show the distribution of change in peptide phosphorylation for PTK, Lep-OLA vs Lep-Veh.

**Supplementary Figure 6.**
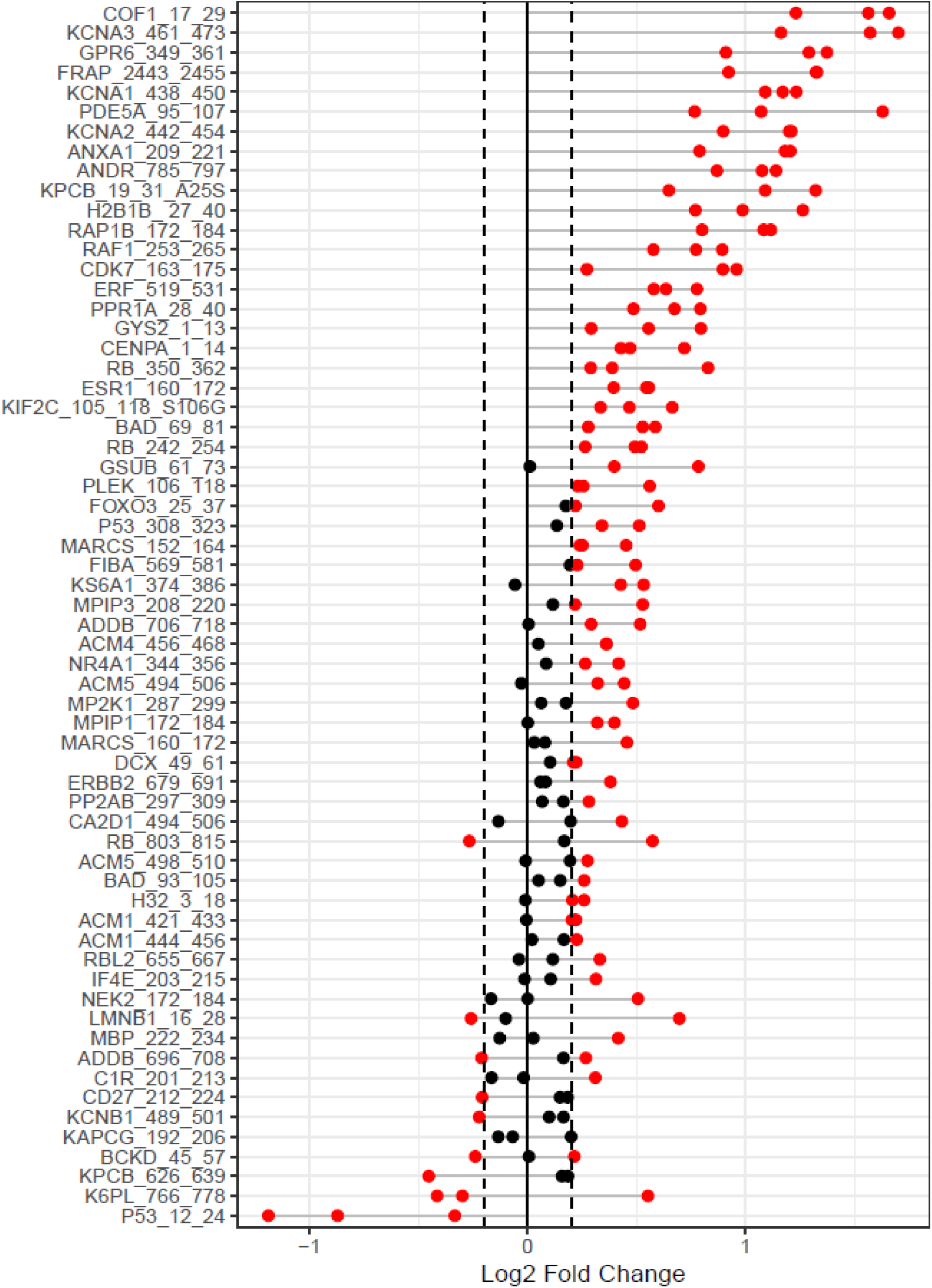
Waterfall Plot to show the distribution of change in peptide phosphorylation for STK, Veh-OLA vs Veh-Veh.

**Supplementary Figure 7.**
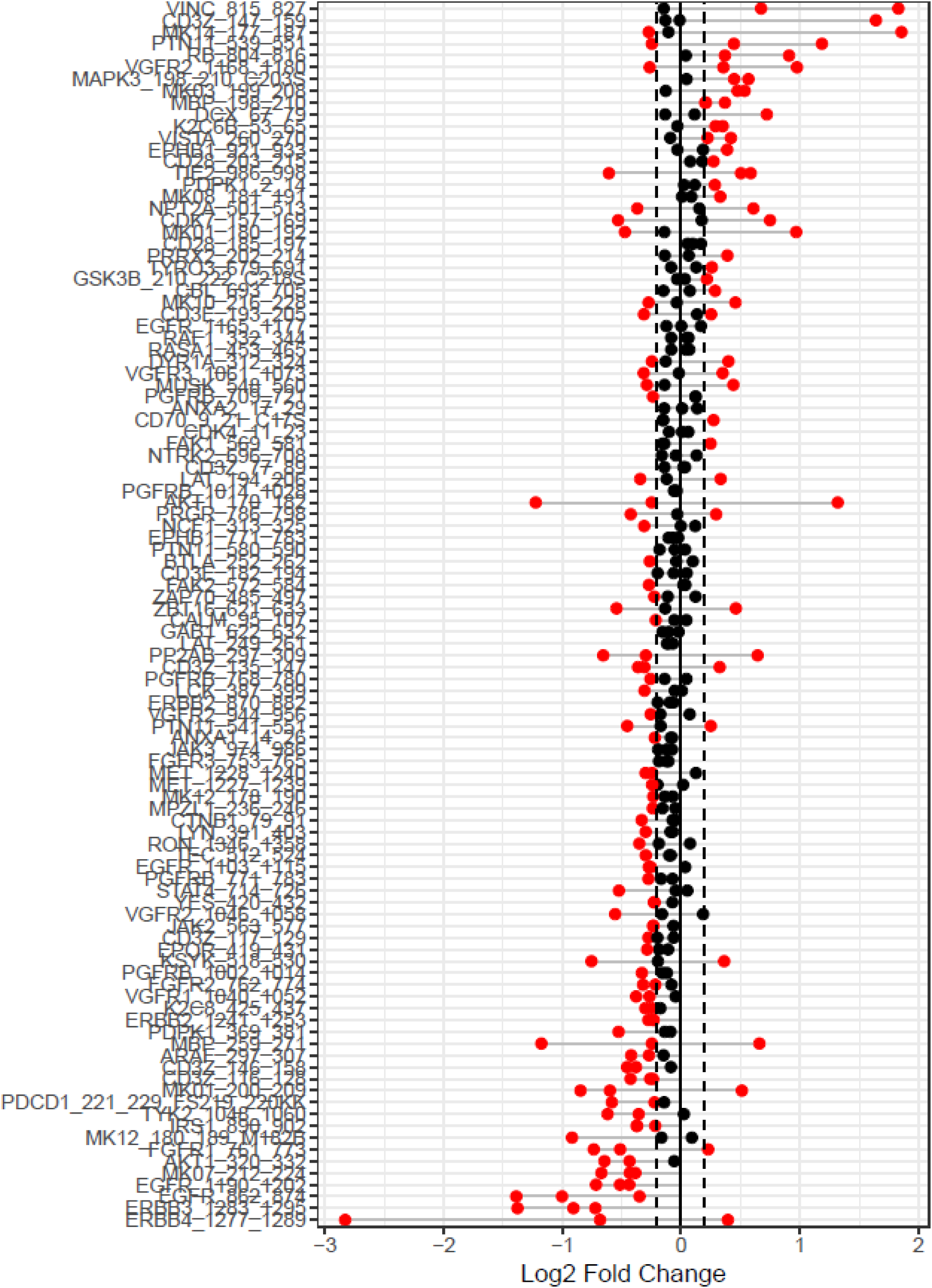
Waterfall Plot to show the distribution of change in peptide phosphorylation for PTK, Yeh-OLA vs. Yeh-Yeh.

